# Non-destructive real-time monitoring of underground root development with distributed fiber optic sensing

**DOI:** 10.1101/2023.07.03.547481

**Authors:** Mika Tei, Fumiyuki Soma, Ettore Barbieri, Yusaku Uga, Yosuke Kawahito

## Abstract

Crop genetic engineering for better root systems can offer practical solutions for food security and carbon sequestration; however, soil layers prevent direct visualization. Here, we demonstrate an original device with a distributed fiber-optic sensor for fully automated, real-time monitoring of underground root development. We demonstrate that spatially encoding an optical fiber with a flexible and durable polymer film in a spiral pattern can significantly enhance sensor detection. After signal processing, the resulting device can detect the penetration of a submillimeter-diameter object in the soil, indicating more than a magnitude higher spatiotemporal resolution than previously reported with underground monitoring techniques. We also developed computational models to visualize the roots of root crops and monocotyledons, and then applied them to radish and rice to compare the results with those of X-ray computed tomography. The device’s groundbreaking sensitivity and spatiotemporal resolution enable seamless and laborless phenotyping of root systems that are otherwise invisible underground.

## 1 Introduction

The root system architecture (RSA) has important implications for food security, climate action, and bio-inspired civil engineering technologies. For food security, the RSA determines crop resilience to environmental stress [1] and nutrient uptake efficiency [2, 3]. With climate change, roots and root exudates are natural solutions for carbon sequestration [4]. Deep and densely rooted plants are advantageous for a larger capacity and stability of carbon storage [5–7]. In civil engineering, biological strategies for excavation, soil penetration, and anchoring can inspire solutions to the geotechnical challenges of unsustainable, energy-intensive processes [8]. Elucidating underground RSA can offer relevant solutions for today’s global issues; however, layers of opaque soil hinder easy visualization.

Underground roots have been studied either destructively or in artificial environments. In destructive methods, root samples are usually excavated by shoveling [9], coring [10], trenching [11], or gravitation [12], then washed and examined. For example, shovelomics combined with laser ablation tomography is used to anatomically characterize nodal roots [13], and backhoe-assisted monoliths combined with RNA-seq are used to reveal the transcriptomic profiles of rice roots [14]. Destructive methods can bring roots under direct vision but are unsuitable for high-throughput and dynamic experiments because of their single-time and laborious measurement. In an artificial system, the surrounding environment is modified to circumvent the light attenuation of the soil using transparent soil substitutes [15, 16], thin chambers [17–19], and turn tables [20–23]. These methods may enable high-throughput and semi-automated measurements but may not be applicable to plants *in situ*.

Several *in-situ* monitoring methods have been developed, such as acoustic imaging [24], electron impedance tomography [25, 26], ground-penetrating radars [27–29], and water potential sensor arrays [12]; however, they have yet to achieve sufficient spatial resolution and sensitivity required for structures as fine as crop roots. The smallest objects detected using these methods were tree roots with diameters of 5 mm for acoustic imaging [24] and 2.5 mm for GPR [27]. The more direct method of inserting a glass tube and camera into the soil has been used to observe fine roots [30, 31]; however, this method is limited in the field of view.

Therefore, current underground visualization techniques lack spatiotemporal resolution, sensitivity, or range for real-time monitoring of RSA.

To achieve requisite spatiotemporal resolution and range, distributed fiber optic sensor (FOS) based on Rayleigh scattering is an emerging technology for monitoring physical quantities such as temperature, strain, and vibration [32]. Fiber optics, which are made of transparent amorphous solids, such as glass and polymers, are widely used in telecommunications. Light waves are trapped in the higher-refractive-index core interfaced with the lower-refractive-index cladding. In contrast to telecommunications, which relies on transmission, FOS exploits the scattering caused by heterogeneity in the core material. Because Rayleigh scattering is material-specific and independent of external energy transfer, the longitudinal coordinates of the FOS can be deciphered using tunable laser-based optical frequency-domain reflectometry [33]. When the fiber is locally stretched by mechanical or thermal expansion, light reflection is delayed, which manifests as an induced shift in the reflected spectrum. The degree of expansion is linearly related to the spectral shift of the segment corresponding to the reference site [34]. FOS has been applied to study large-scale dynamics such as earthquakes [35], glacial flows [36], and whales [37]. Innovation is required for its application in root development, which operates many orders of magnitude smaller in force and energy.

The objective of this study was to overcome the difficulties related to the sensitivity and spatial resolution of FOS required for monitoring root development. To do so, we designed and implemented an original device that significantly enhanced the FOS sensitivity and spatial encoding, accompanied by signal processing and visualization software for RSA determination. The completed device and software were validated by real-time root monitoring for radish and rice over weeks and months, and the virtually reconstructed root structures generally agreed with high-resolution X-ray computed tomography (CT) images of the actual roots.

## 2 Results

### 2.1 Design and characterization of the sensing device

The proposed device is a spatially encoded FOS connected to a commercially available optical reflectometer. Fig. 1 shows the complete design of the device. The FOS consisted of a single-mode fiber composed of a GeO_2_-doped silica core, pure silica cladding, and a polyimide coating. To cover the space for the entire root system, the FOS is supported by rigid structures that are extensively and evenly placed in the soil. The combination of structuring and optical frequency-domain reflectometry enables the spatial encoding of one-dimensional signals. FOS density of 1 per 30 mm is attempted for detecting root diameter of 0.3 mm for 15 µ_ε_ [21] according to Eq. A1 [34].

**Fig. 1.**
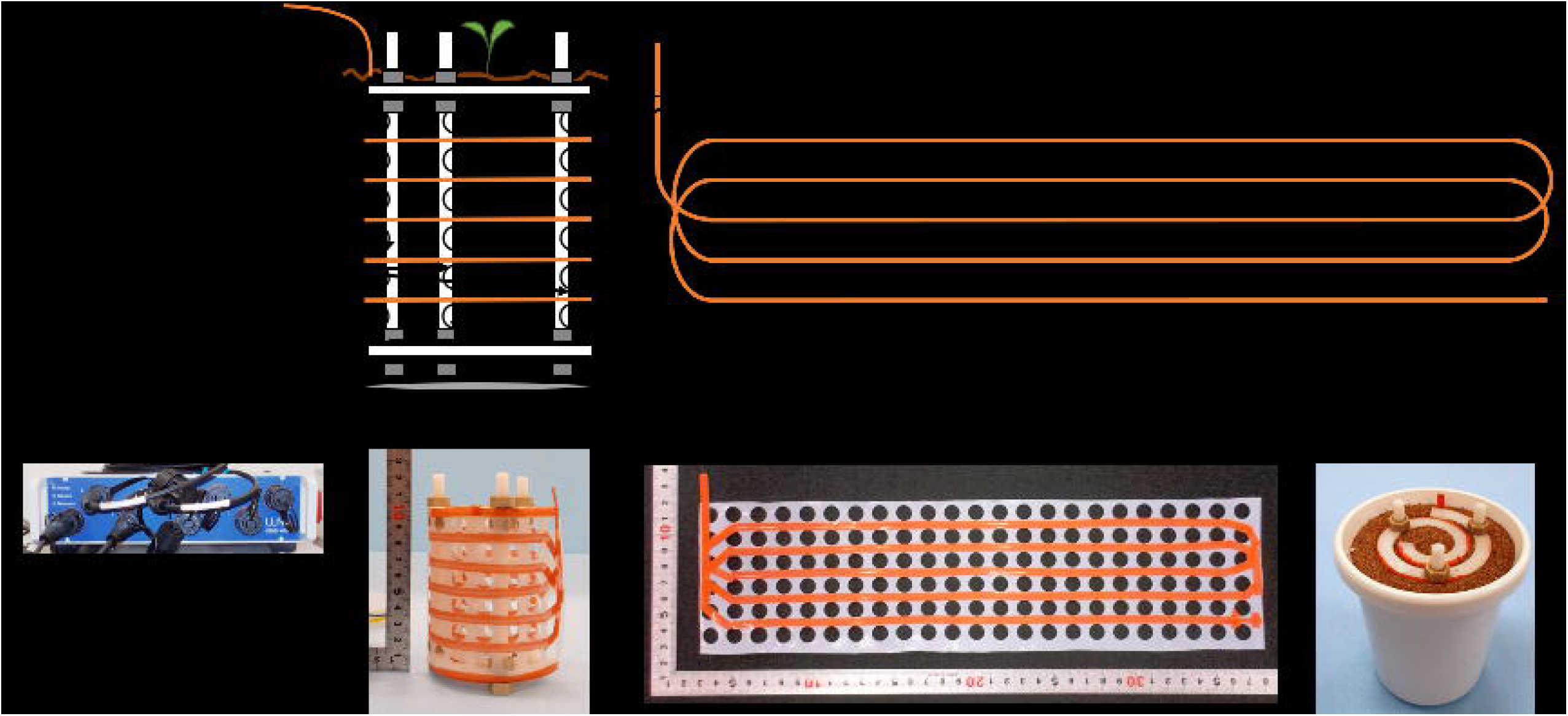
A schematic and photos of the completed FOS device. (a) The overall sensing schemes. FOS is attached to a perforated polymer film which is mechanically supported. (b) The unfold schematic of the device. (c) A photo of the reflectometer. (d) The side view of the device. (e) The unfold view of the device. (f) The device filled with Profile in a cultivation pot.

Distributed strains were recorded to evaluate the sensitivity of the device, while a thin metal wire mimicking the root penetrated the center. The direct installation of the FOS results in a low signal and high noise, as shown in Fig. S1. To enhance the signal gain and stability, the FOS was attached to a perforated polymer film to increase the surface area and plasticity of the sensor. Polyoxymethylene (POM) and polytetrafluoroethylene (PTFE) with Young’s moduli of 3.015 and 0.569 GPa, respectively, were used in the prototype, as shown in Fig. S2. As expected, both polymers improved the gain (stress-to-strain ratio) and stability, with PTFE demonstrating superior performance.

Different FOS orientations were tested on the PTFE films, as shown in Fig. 2. The horizontally and vertically oriented FOS on PTFE exhibited high sensitivity and stability. However, the linearity was compromised in the vertical orientation as the strain saturated at 1 MPa compression of the filler by the wire. Different penetration sizes and media stiffnesses were tested using agarose gel to fill the cultivation pot with horizontally fixed FOS on the PTFE film, and the results are plotted in Fig. S3. The force required for penetration, as well as the strain, increased as the size of the penetrating object increased or at a higher concentration of agarose. The amplitude of strains showed proportionality to the force of penetration, with a detection limit of ¡ 0.07 N when a metal wire of 0.5 mm diameter was penetrated in 1.5 % agarose gel. This suggests that our device is highly sensitive to the force exerted by the root development, regardless of the media material.

**Fig. 2.**
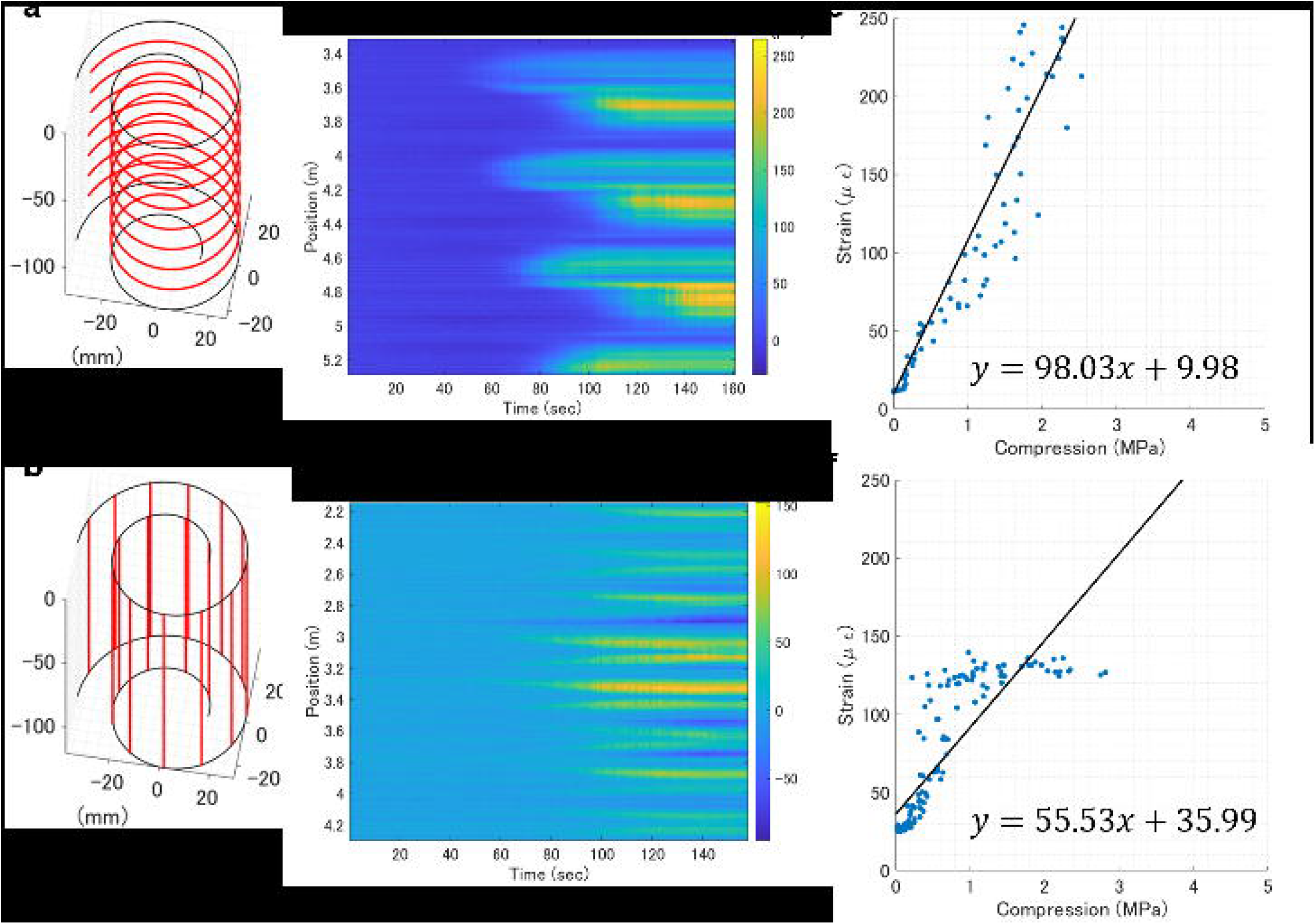
Evaluation of FOS orientation. (a, b) FOS placement of a prototype indicated by red lines. (c, d) Distributed strain recording from a horizontally or vertically placed FOS when penetrating 1.2 mm-diameter metal wire at 1 mm/sec into the glass beads filling the device and the surrounding. (e, f) Stress-strain curves of (c) and (d), respectively.

Therefore, the complete design of the sensing device consisted of an FOS horizontally fixed on a perforated PTFE film. The intervals between adjacent segments of the FOS were in the range of 15–30 mm. The size and frequency of the perforations were adjusted to reduce interference on root development in a trade-off with signal sensitivity and stability.

### 2.2 Noise reduction and temperature separation

The FOS was sealed between chemically stable PTFE films within the device to avoid non-target chemical-based effects on the optical signals [38].

Digital signal processing was also used to reduce noise. Inherent noise from the reflectometry was removed using a Butterworth low-pass or median filter in the spatial domain. Smoothing is also necessary for the subsequent root visualization of rice plants, in which peak finding is used to estimate the locations of root tips. In parallel, notch filters with cutoff frequencies of approximately 1/day were applied to separate the background noise caused by the diurnal temperature effect. Residual transient noise was removed by elementary subtraction of the spatial averages. The background physical strain caused by the spiral structure was compensated for by the elementary subtraction of the temporal averages for the duration before germination, for example, the first 24 h of cultivation.

Fig. 3 shows an example of distributed strain recording for root growth in a radish. Notch filters and elementary subtraction of spatial averaging separate the ambient temperature contributions. The separated signals closely followed the temperature of the growth chamber; the averaged signals from the above-ground portion of the FOS were nearly identical to the ambient temperature, whereas the average underground signals had smaller amplitudes and delays of up to 2.2 hours (Fig. 3a). The smaller amplitudes and delays in the temperature can be attributed to the insulation and latent heat of the soil.

**Fig. 3.**
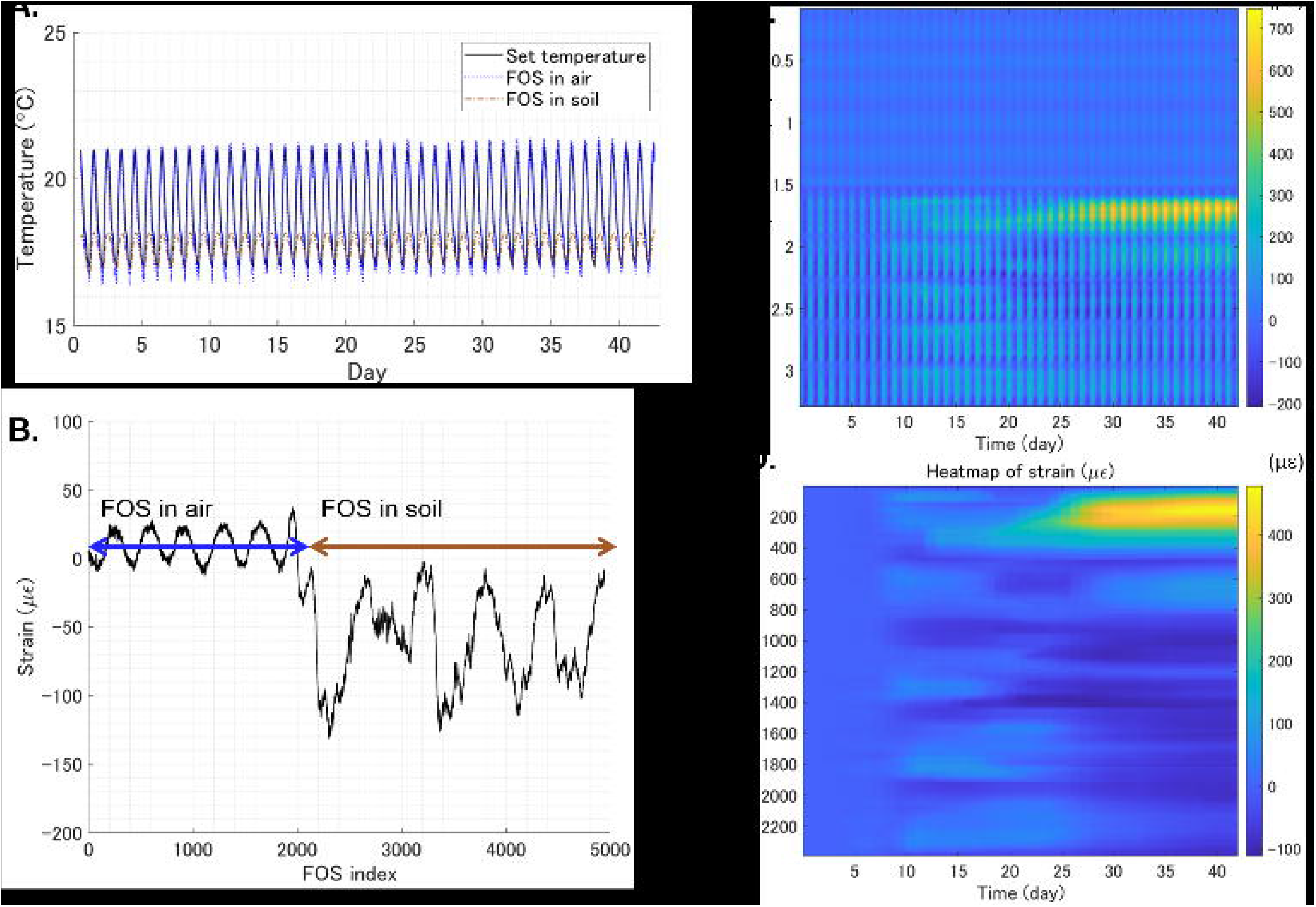
Digital signal processing on a sample data recording radish growth. (a) Comparison of time-series FOS signal averages and growth chamber temperatures. (b) The background physical strain indicated by the FOS signal averages for the pre-germination period. (c) Distributed strain recording for the full FOS length before signal processing. (d) Distributed strain recording for the underground portion of FOS after signal processing.

Raw signals from FOS in soil should be adjusted for the thermal expansion of PT FE (125 µ_ε_ °C*^−^*^1^), which is approximately 220 times larger than silica (0.55 µ_ε_◦°C*^−^*^1^). The structural bias obtained by temporal averaging indicated two portions of the FOS: a loose bare fiber reeled above the ground and a tightly strained fiber fixed onto the PTFE underground (Fig. 3b).

As the radish roots grew, local accumulation of the strains was observed (Fig. 3c-d). Because a slight periodic background is still apparent on the timescale, a mathematical morphology filter can substitute frequency filters for better results. Mathematical morphology filters (MMF) differentiate based on a waveform and can thus remove noise in the overlapping frequencies with the target signals [39][40]. The MMF for a triangular waveform, instead of notch filters, effectively removed the temporal noise, as shown in Fig. S4.

### 2.3 Virtual reconstruction of radish root

To visualize the root development in radishes, we reconstructed a computational model for secondary root growth. Root development has two phases: primary growth occurs as a vertical elongation, and secondary growth occurs as a lateral expansion of the root volume. As the details of the secondary root growth model are described in Supplementary Theory I, the model assumes that FOS deformation is caused by root volume expansion and estimates the root radius from distributed strain measurements according to Eq. B2.

The supplementary movie depicts the time-series distributed strain measurement and virtual reconstruction of the radish root over 42 days after sowing. At the end of the cultivation, an X-ray CT scan and image reconstruction were performed, as shown in Fig. 4. Because the implementation of the FOS limits the vertical resolution to >15 mm, the reconstruction from the distributed strain is coarser than that from the X-ray CT. Nevertheless, the depth of root growth was consistent between the two visualization methods. Dynamic tracking by FOS indicat−ed that the root laterally expanded near the surface on day 12, followed by mild expansion leftward from day 18 to 25, and rapid expansion from day 26 to 37. A localized increase in the distributed strain indicated root growth at z = 60 mm on day 22, but the root was hardly visible in the X-ray CT images. The radish root excavated at z = 60 mm had a diameter of 0.8 mm (Fig. S4). This suggests that our device can detect a root as fine as 0.4 mm in radius, similar to the standard X-ray CT limit for root detection without image processing [21].

**Fig. 4.**
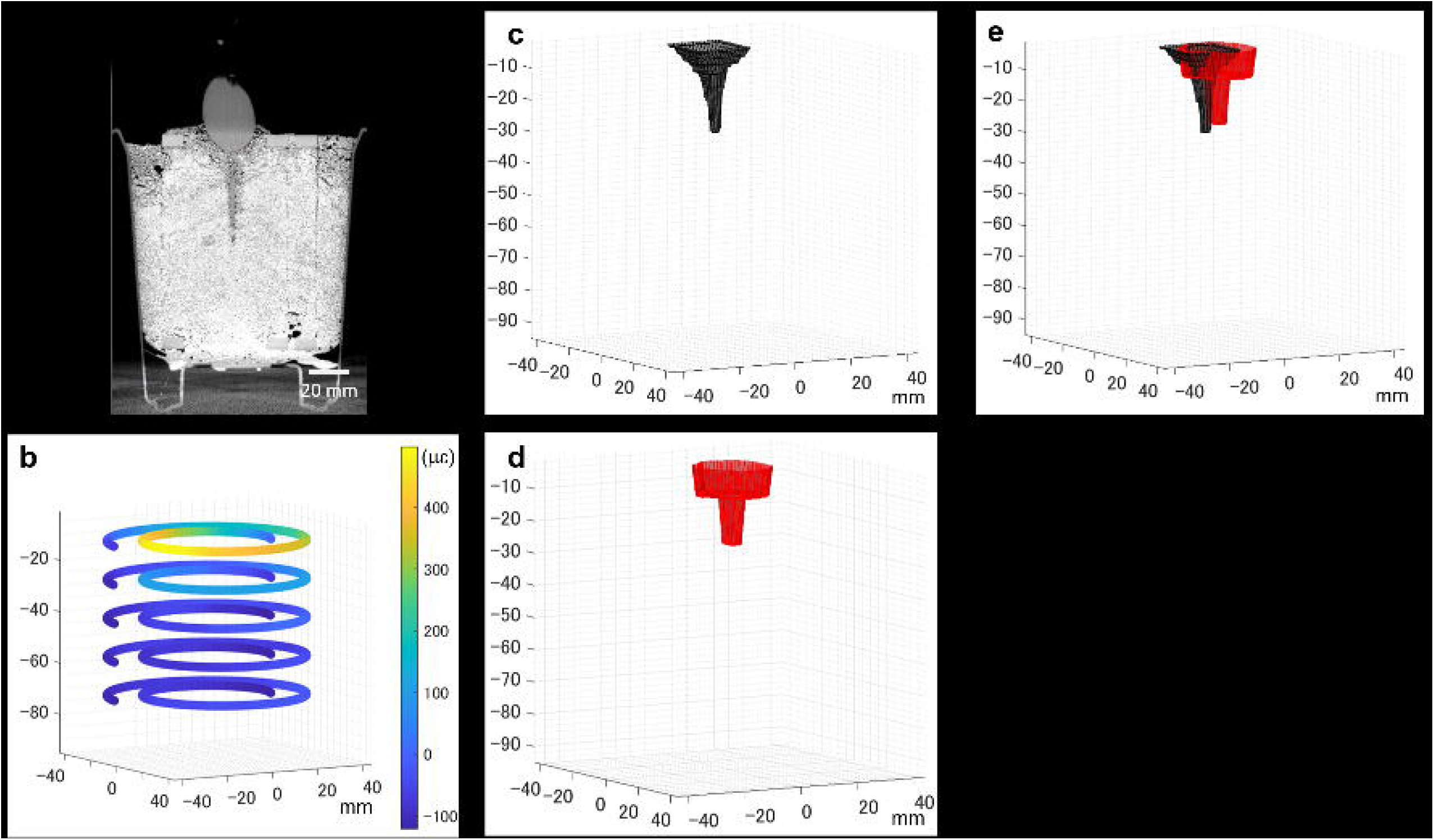
Visualization results from t h e X-ray CT and FOS measurement on the 42nd day of radish cultivation. (a) An image slice from the X-ray CT. (b) The spatially encoded heatmap indicating distributed strain and its 3D coordinate from FOS measurement. (c) Virtual radish reconstruction from t h e X-ray CT images by drawing a surface for the root-soil border. (d) The virtual radish reconstruction from the FOS measurement, according to Eq. A1. (e) Superimposition of panels c and d.

### 2.4 Virtual reconstruction of rice roots

To visualize the RSA of the rice plants, we reconstructed a computational model for primary root growth using distributed strain measurements. The details of the model are provided in Supplementary Theory II. Unlike the radish model, the rice model assumes that the strains are caused by the downward force exerted at the root tip and estimates the 3D coordinates of the force source. The analytical framework can only deduce the distance between the sensor and the root tip. An optimization framework was used to determine the 3D coordinates of the root tips. First, peak-finding was used to initialize the coordinates for all root tips. After spatial low-pass filtering, the location of the FOS index with the local maximal strain was assumed to be the root tip. Then the least-squared-errors optimization is used on a root tip to match the array of simulated strains ε_θθ_ from Eq. B5 to the array of the actual measurements for every FOS index.

Finally, the base of the stem and coordinates of the root tip are connected by a hyperbolic line to reconstruct the entire root.

Distributed strain recordings from different rice cultivation pots are shown in Fig. S5. Root growth can generally be detected as an increase in local strains starting from day 10 to 13 after sowing. Virtual rice roots were reconstructed at a few time points in which cultivation was disrupted to obtain X-ray CT scans, and the two visualization methods at the same time point are compared in Fig. 5. Both methods revealed similar root structures when the crown roots grew within the inner layer of the spiral (Fig. 5a-c). Initialization fails when the crown roots are underdeveloped, whereas the primary root penetrates the center of the spiral (Fig. 5d). Conversely, when crown roots spread widely beyond the inner spiral layer, the FOS senses negative strains due to forces from the opposite side (Fig. 5e-f). Negative strains were either rejected by the threshold (Fig. 5e) or interfered with the other root tips that developed inside the inner spiral layer (Fig. 5f) during reconstruction.

**Fig. 5.**
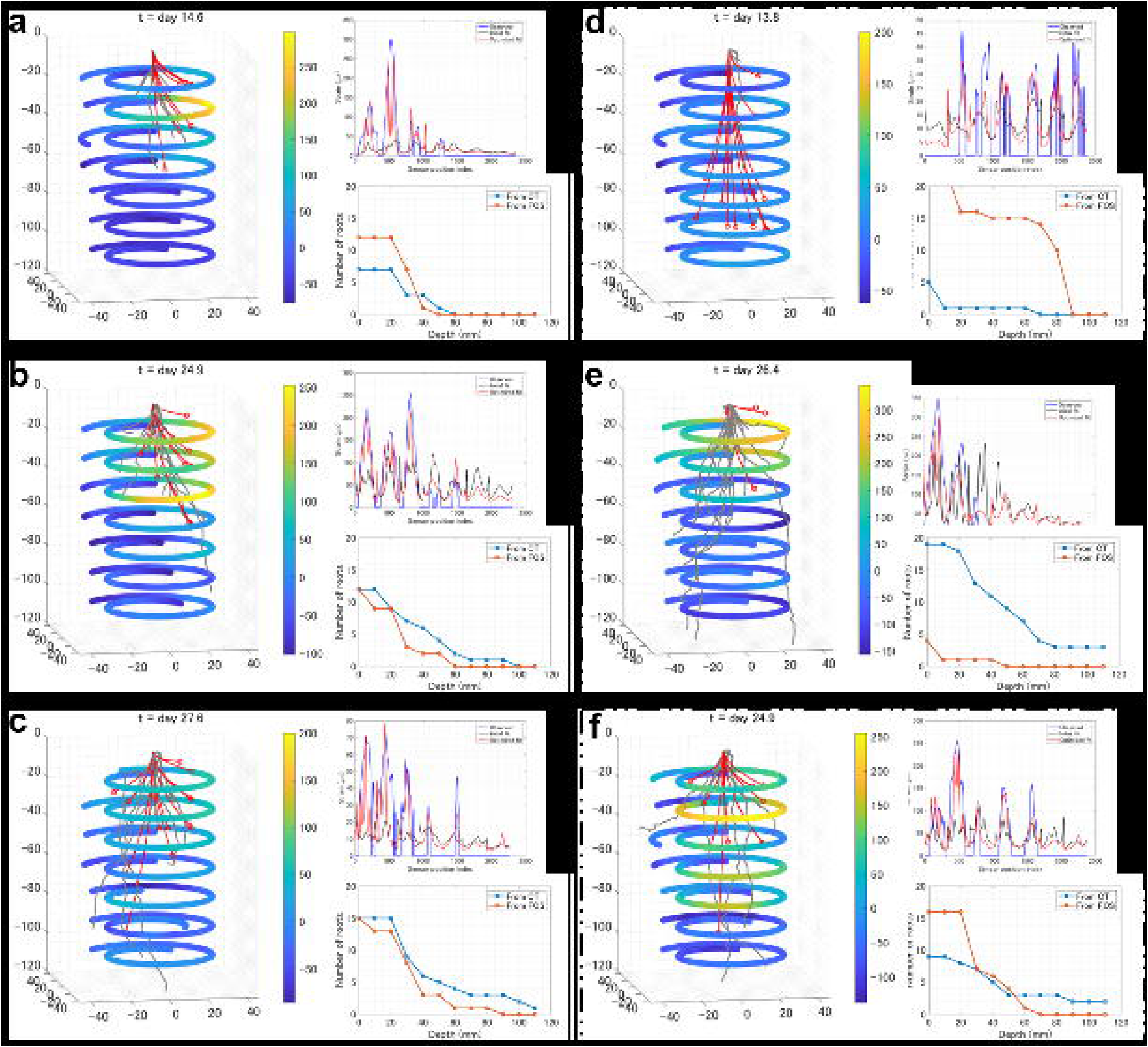
Visualization results from the X-ray CT and FOS measurement of several rice cultivation pots at different growth stages. (a) (left) Superimpositions of data obtained on day 15 for Pot #2. Spatially encoded strain heatmap is overlayed with reconstructed rice roots; gray lines indicate CT reconstruction, and red lines indicate FOS reconstructions. (top right) Plots of distributed strain measurement from FOS and simulated strains from the computational model. (bottom right) Plots counting the number of roots along depth for CT and FOS. (b) Data for Pot #6 on day 25. (c) Data for Pot #7 on day 28. (d) Data for Pot #4 on day 14. (e) Data for Pot #1 on day 27. (f) Data for Pot #4 on day 25.

## 3 Discussion

The FOS is a promising real-time monitoring technology that is easy to install with a submillimeter longitudinal resolution and is suitable for long-term experiments. We developed a device that enhances FOS sensing and computational abilities to monitor underground root development. We named the device “Fiber-RADGET” for “Fiber optic sensor-based RADicle gadGET.” The Fiber-RADGET prototype explored different materials and orientations of the device backbone. FOS sensitivity is greatly improved by immobilizing FOS on polymer films such as POM and PTFE. We designed and implemented a spiral concentric backbone that encased the target object to provide spatial encoding for the FOS. The horizontal orientation of the FOS structured on the PTFE film was the most effective for detecting a thin metal wire that mimicked a crop root. The completed Fiber-RADGET design was used to track root development over weeks and months. The sensing mechanism relies on the local strain increase caused by nearby root growth. Computational models were developed for the root visualization of radish and rice. The virtually reconstructed roots from the FOS generally concorded with the roots traced in high-resolution X-ray CT scans of the actual roots, except in cases with underdeveloped roots or roots grown behind the polymer film.

Although radish and rice have two distinct root structures, Fiber-RADGET detected the root growth in both, as a local strain that increased with FOS. For the software, we developed two different computational models focusing on volume and pressure to visualize more types of roots, covering root crops and monocots. Root crops are convenient to model because an enlarged root induces significant deformation in the FOS. However, to apply the FOS to general dicots other than root crops, such as radish, the model should distinguish the two deformation patterns caused by the primary and lateral roots.

The accuracy of our pressure-focused primary root growth model was acceptable, despite multiple simplifications. One assumption is that the force exerted by the root tip is coaxial with gravity. This assumption was based on root gravitropism [41]. Although this applies to most existing cases, crop genetic engineering will likely target the genes responsible for root growth in the direction of optimizing the RSA for water and nutrient uptake [3]. Therefore, the assumption of gravitropism must be re-confirmed in these plants. If necessary, an increased degree of freedom for the pressure source should be considered in the model, combined with better heuristic or deep-learning data for more accurate reconstruction.

To date, no high-resolution nondestructive visualization methods have been developed for crop roots in open fields. The Fiber-RADGET should be swiftly translatable to fields like other FOS applications [32, 35, 37, 42]. Extensive characterization of environmental signals is required *a priori* to decouple the root development signals from compound measurements for accurate visualization. On the other hand, by decoupling signals, Fiber-RADGET can be a generic environmental monitor that may simultaneously track temperature, soil compaction, and other biological activities. For example, air and soil temperatures were extracted from background signals in our radish demonstration. With appropriate signal processing using bandpass filters and cross-correlation, many signals must be isolated and tracked separately to reveal the concealed underground dynamics.

With further adjustments in the device and software, Fiber-RADGET can be applied in open fields, enabling real-time monitoring of field crop roots.

## 4 Methods

### 4.1 Instrumentation

A high-definition fiber optic strain sensor, HD6S (Luna Inc., Virginia, USA), was used as the FOS., 8-channel ODiSI 6100 reflectometer (Luna Inc., Virginia, USA) was used for high-throughput recording. 0.1-0.2 mm thick NITOFLON (Nitto Denko, Osaka, Japan) and 0.2 mm thick polyacetal (Misumi Group, Tokyo, Japan) sheets were used as polymer films in the prototypes. Perforation was performed using a film punch press UDP5000 (Fuji Shoko Machinery, Saitama, Japan) with circular holes 5 or 10 mm in diameter at intervals of 15 mm. The FOS and film were fixed by 5 mm wide 0.1 mm thick NITOFLON tape. The backbone structure was built using polyamide screws (Esco, Osaka, Japan), polycarbonate hexagonal nuts, and 5 mm thick polyacetal plates (Esco). For the completed design, the plates were cut into spiral shapes using a laser cutter Speedy 300 (Trotec, Marchtrenk, Austria). The polymer film is fixed along a pair of spiral plates. The number of turns of the spiral is 1.75 turn in Fig. 1, 3, and 4, for wider pot applications and 1.5, as shown in Fig. 2, Fig. S2, Fig. S3 for narrower-pot applications. For the direct installation of the FOS in Fig. S1, a PTFE tube with 0.3 mm diameter (Hagitech, Chiba, Japan) was used instead of a film to protect the FOS, and three concentric circles were used to hold the tube. The structured FOS was placed in an empty pot and filled with materials such as gel, beads, and soil. For stress-strain analysis, strains were recorded twice every second. For plant growth, strains were recorded twice every 10 min.

### 4.2 Stress-strain analysis

For abiotic characterization of FOS with stress-strain analysis, stainless wires of 0.5-1.2 mm diameter (Hikari, Osaka, Japan) or the flat side of the bamboo skewer of 2.5 mm diameter (Izumo Chikuzai Industry, Shimane, Japan) were used as the stress input. For filling, Profile Greens Grade (Profile Product, IL, USA) was used to test for the direct installment of FOS and compare effects from different polymer films, globular glass beads FGB20 (Fuji Manufacturing; Tokyo, Japan) of 0.71-1000 mm diameter were used to compare FOS orientations, and agarose gel in tris/acetic acid/EDTA (Bio-rad, California, USA) were used for different stiffness. Force tester MCT1150 (A&D, Tokyo, Japan) was used to measure the force and pressure during wire penetration.

### 4.3 Digital signal processing

Noises in strain recording were removed using either the combination of frequency filters or MMF. The noise reduction based on frequency was obtained by inverting the pointwise product of the Fourier transform of the strain recording and the superposition of the notch filter with the stopband of (duration of the experiment)/ (24 hour) and a band-pass filter with the cutoff of (length of the FOS)/ (30 mm). The noise reduction based on mathematical morphology was obtained by sequentially averaging the closing of the opening and opening of the closing of the strain recording with the triangular structures of the varying size, according to the implementation described in [39]. The largest structure used was the isosceles triangle with a base of 2.5 days. For the background compensation, the average over time was subtracted for every position and the mean of FOS was subtracted at every timepoint from the strain recording.

### 4.4 Plant materials

The radish (*Raphanus raphanistrum*) line of Kouhaku (Sakata Seed, Kanagawa, Japan) and rice (*Oryza sativa*) lines of Kinandang Patong (KP, IRGC#23364) were used as sample crops in this study. Radish seeds were washed with 70 % ethanol for 20 s, rinsed three times with autoclaved water, and stirred in a 2.5 % hypochlorite solution for 15 min before being rinsed and incubated with autoclaved water for germination. The KP seeds were immersed in 0.25 % Techleed C Flowable (5 % ipconazol, 4.6 % Cu(OH)_2_; Kumiai Chemical Industry; Tokyo, Japan) at 15°C for 24 hours and in water at 30°C for two days. Each germinated seed was sown at the center of the soil at shallow depth. The combination of the FOS and plant materials information (dates of sowing and ending experiments) is given in Table 1.

**Table 1.** Pot number description

| Pot # | FOS ID | Start date | End date |
| --- | --- | --- | --- |
| 1 | FS2020LUNA063319 | July 1st, 2021 | July 28th, 2021 |
| 2 | FS2020LUNA063328 | July 1st, 2021 | July 28th, 2021 |
| 3 | FS2020LUNA063333 | July 1st, 2021 | July 28th, 2021 |
| 4 | FS2020LUNA063328 | August 6th, 2021 | September 10, 2021 |
| 5 | FS2020LUNA063333 | August 6th, 2021 | September 10, 2021 |
| 6 | FS2020LUNA063338 | August 6th, 2021 | September 10, 2021 |
| 7 | FS2020LUNA063339 | August 6th, 2021 | September 10, 2021 |

### 4.5 Growth conditions

To use Profile as soil, it was rinsed with tap water three to five times and dried before being filled into four cultivation pots, 70 mm in diameter and 150 mm in height. The profile was saturated pre-cultivation, with a modified Kimura B solution (1.23 mM KNO3, 1 mM KCl, 0.37 mM CaCl_2_, 0.55 mM MgSO_4_, 0.18 mM KH_2_O_4_, and 8.9 µM Fe(II)-EDTA (pH 5.5)) For rice, the saucer was filled with the modified Kimura B solution to a depth of 15 mm. For the radishes, the saucer was filled with tap water to a depth of 30 mm. The cultivation was conducted using Biotron (Nippon Medical & Chemical Instruments; Osaka, Japan) with the conditions following the 24-h cycle emulating an average day of Kanagawa in May 2021 (17–21 °C) or Tsukuba, Japan in July 2017 (25-30 °C, 0-0.5 mmol photosynthetic photon/m^2^/s), for radish and rice, respectively. The humidity was set to 50 % during the day and 60 % at night. The CO2 level remained at an average of 400–500 ppm.

### 4.6 Root visualization

Details about plant root reconstruction from FOS are provided in the Supplementary Theories. The model parameters and optimization results for the primary root growth model are listed in Table 2. The custom code for the virtual root reconstruction in MATLAB (MathWorks, Massachusetts, USA) is available at https://github.com/mtei1/Fiber-RADGET.git. X-ray CT scans were obtained approximately 14 and 28 d after sowing using the X-ray CT system InspeXio SMX-225CT FPD HR (Shimadzu Corporation, Kyoto, Japan), as previously reported [43]. The coordinates for the radish boundaries were manually extracted using Fiji [44], whereas RSAtrace3D in combination with RSAvis3D was used to extract the rice root coordinates [21, 45].

**Table 2.** Parameters used for rice reconstruction.

| Pot # | Day after sowing | $P$ (GPa) | Initial<sup>1</sup> | Optimal<sup>2</sup> |
| --- | --- | --- | --- | --- |
| 1 | 15 | 1e5 | 0.8015 | 0.6407 |
| 2 | 15 | 5e5 | 0.7586 | 0.5429 |
| 3 | 15 | 2e5 | 0.8413 | 0.6654 |
| 1 | 27 | 2.5e6 | 0.865 | 0.5826 |
| 2 | 27 | 5e5 | 0.9734 | 0.6664 |
| 3 | 27 | 2e5 | 0.8812 | 0.7108 |
| 4 | 25 | 4e5 | 0.7483 | 0.447 |
| 5 | 25 | 2e5 | 0.8415 | 0.6372 |
| 6 | 25 | 6e5 | 0.7312 | 0.4341 |
| 7 | 25 | 3e5 | 0.7013 | 0.5054 |
| 4 | 28 | 2e5 | 0.911 | 0.7106 |
| 5 | 28 | 1e5 | 0.8512 | 0.6555 |
| 6 | 28 | 2e5 | 0.9486 | 0.6375 |
| 7 | 28 | 1e5 | 0.817 | 0.5878 |

$G = 0.1$ (GPa) and $\nu = 0.5$ for all pot numbers.
<sup>1</sup>Normalized residual sum of square at initialization.
<sup>2</sup>Normalized residual sum of square after optimization.

## Supplementary information

This article has an accompanying supplementary movie file.

## Author contributions

MT, YK, and YU conceived of and designed the experiments. MT conducted instrumentation and abiotic experiments and analyzed all data. MT and FS conducted the plant experiments and recorded the data. MT and EB constructed the computational models and simulated them for reconstruction. All the authors contributed to the manuscript.

## Acknowledgments

This work was supported by the Cabinet Office, Government of Japan, Moonshot R&D Program for Agriculture, Forestry, and Fisheries (funding agency: Bio-oriented Technology Research Advancement Institution), Japan Science and Technology Agency (JST) CREST Grant Numbers JPM JCR17 O1 and JPM JCR17 O3, and MEXT/JSPS KAKENHI. Grant Number: JP21K14779. We thank Mr. Yuji Manabe of the Japan Agency for Marine-Earth Science and Technology for their limitless assistance in conducting this research. We also thank Dr. Saki Yoshida at the Tokyo University of Agriculture for their advice in assessing the preliminary data, and Dr. Shota Teramoto and Ms. Yoko Fukuda for assistance with the X-ray CT.

## References

[1] Mickelbart, M.V., Hasegawa, P.M., Bailey-Serres, J.: Genetic mechanisms of abiotic stress tolerance that translate to crop yield stability. Nat. Rev. Genet 16(4), 237–251 (2015)

[2] Gruber, B.D., Giehl, R.F., Friedel, S., von Wirén, N.: Plasticity of the arabidopsis root system under nutrient deficiencies. Plant Physiology 163, 161– 179 (2013)

[3] Yamazaki, K., Fujiwara, T.: The effect of phosphate on the activity and sensitivity of nutritropism toward ammonium in rice roots. Plants 11(6) (2022).

[4] Lal, R.: Carbon sequestration. Phil. Trans. R. Soc. B 363, 815–830 (2008)

[5] Yang, Y., Tilman, D., Furey, G., Lehman, C.: Soil carbon sequestration accelerated by restoration of grassland biodiversity. Nat. Comm. 10, 718 (2019)

[6] Button, E.S., et al.: Deep-c storage: Biological, chemical and physical strategies to enhance carbon stocks in agricultural subsoils. Soil Biology and Biochemistry 170, 108697 (2022).

[7] Eckardt, N.A., et al.: Climate change challenges, plant science solutions. The Plant Cell 35(1), 24–66 (2022)

[8] Martinez, A., et al.: Bio-inspired geotechnical engineering: principles, current work, opportunities and challenges. Géotechnique 72(8), 687–705 (2022).

[9] Trachsel, S., Kaeppler, S.M., Brown, K.M., Lynch, J.P.: Shovelomics: high throughput phenotyping of maize (zea mays l.) root architecture in the field. Plant and Soil 341, 75–87 (2011)

[10] Teramoto, S., Kitomi, Y., Nishijima, R., Takayasu, S., Maruyama, N., Uga, Y.: Backhoe-assisted monolith method for plant root phenotyping under upland conditions. Breeding Science 69, 508–513 (2019).

[11] Teramoto, S., Uga, U.: A deep learning-based phenotypic analysis of rice root distribution from field images. Plant Phenomics, 3194308 (2020).

[12] Dowd, T.G., Li, M., Bagnall, G.C., Johnston, A., Topp, C.N.: Root System Architecture and Environmental Flux Analysis in Mature Crops using 3D Root Mesocosms.

[13] Oyiga, B.C., et al.: Genetic components of root architecture and anatomy adjustments to water-deficit stress in spring barley. Plant Cell Environ. 43, 692–711 (2020)

[14] Kawakatsu, T., et al.: The transcriptomic landscapes of rice cultivars with diverse root system architectures grown in upland field conditions. Plant J. 106, 1172–1190 (2021)

[15] Adachi, H., Ozawa, M., Yagi, S., Seita, M., Kondo, S.: Pivot burrowing of scarab beetle (trypoxylus dichotomus) larva. Sci. Rep. 11, 14594 (2021)

[16] Ma, L., et al.: Hydrogel-based transparent soils for root phenotyping in vivo. Proc. Natl. Acad. Sci. U. S. A. 116(22), 11063–11068 (2019)

[17] Wenzel, W.W., Wieshammer, G., Flitz, W.J.: Novel rhizobox design to assess rhizosphere characteristics at high spatial resolution. Plant and Soil 237, 37–45 (2001)

[18] Nagel, K.A., et al.: Growscreen-rhizo is a novel phenotyping robot enabling simultaneous measurements of root and shoot growth for plants grown in soil-filled rhizotrons. Functional Plant Biology 39, 891–904 (2012)

[19] Zengler, K., et al.: Eco-fabs: advancing microbiome science through standardized fabricated ecosystems. Nat. Methods 16, 567–571 (2019)

[20] Heeraman, D., Hopmans, J., Clausnitzer, V.: Three dimensional imaging of plant roots in situ with x-ray computed tomography. Plant and Soil 189, 167– 179 (1997)

[21] Teramoto, S., Takayasu, S., Kitomi, Y., Arai-Sanoh, Y., Tanabata, T., Uga, Y.: High-throughput three-dimensional visualization of root system architecture of rice using x-ray computed tomography. Plant Methods 16, 66 (2020)

[22] Mairhofer, S., Zappala, S., Tracy, S., Sturrock, C., J., B.M., Mooney, S.J., Pridmore, T.P.: Recovering complete plant root system architectures from soil via x-ray µ-computed tomography. Plant Methods 9, 8 (2013)

[23] van Dusschoten, D., et al.: Quantitative 3d analysis of plant roots growing in soil using magnetic resonance imaging. Plant Physiology 170, 1176– 1188 (2016)

[24] Proto, A.R., Iorio, A.D., Abenavoli, L.M., Sorgonà, A.: A sonic root detector for revealing tree coarse root distribution. Sci. Rep. 10, 8075 (2020)

[25] Balwant, P., Jyothi, V., Pujari, P.R., Dhyani, S., et al.: Tree root imaging by electrical resistivity tomography: geophysical tools to improve under-standing of deep root structure and rhizospheric processes. Trop. Ecol. 63, 319–324 (2022).

[26] Corona-Lopez, D.D.J., Sommer, S., Rolfe, S.A., Podd, F., Grieve, B.D.: Electrical impedance tomography as a tool for phenotyping plant roots. Plant Methods 15, 49 (2019)

[27] Yamase, K., et al.: Ground-penetrating radar estimates of tree root diameter and distribution under field conditions. Trees 32, 1657–1668 (2018)

[28] Liu, X., Dong, X., Xue, Q., Leskovar, D.I., Jifon, J., Butnor, J.R., Marek, T.: Ground penetrating radar (gpr) detects fine roots of agricultural crops in the field. Plant and Soil 423(1-2), 517–531 (2018)

[29] Sunvittayakul, P., et al.: Cassava root crown phenotyping using three-dimension (3d) multi-view stereo reconstruction. Sci. Rep. 12, 10030 (2022)

[30] Johnson, M.G., Tingey, D.T., Phillips, D.L., Storm, M.J.: Advancing fine root research with minirhizotrons. Environmental and Experimental Botany 45(3), 263–289 (2001).

[31] Rahman, G., et al.: Soilcam: A fully automated minirhizotron using multispectral imaging for root activity monitoring. Sensors 20(3) (2020).

[32] Soga, K., Luo, L.: Distributed fiber optics sensors for civil engineering infrastructure sensing. J. of Struct. Integrity and Maintenance 3(1), 1–21

[33] Soller, B.J., et al.: High resolution optical frequency domain reflectometry for characterization of components and assemblies. Opt. Express 13, 666– 674 (2005).

[34] Kreger, T., et al.: High Resolution Distributed Strain or Temperature Measurements in Single- and Multi-mode Fiber Using Swept-Wavelength Interferometry. paper ThE42 in Optical Fiber Sensors, OSA Technical Digest (CD) (Optica Publishing Group, 2006) (2006)

[35] Lindsey, N.J., Dawe, T.C., Ajo-Franklin, J.B.: Illuminating seafloor faults and ocean dynamics with dark fiber distributed acoustic sensing. Science 366(6469), 1103–1107 (2019)

[36] Walter, F., et al.: Distributed acoustic sensing of microseismic sources and wave propagation in glaciated terrain. Nat. Commun. 11, 2436 (2020).

[37] Bouffaut, L., et al.: Eavesdropping at the speed of light: Distributed acoustic sensing of baleen whales in the arctic. Frontiers in Marine Science 9 (2022).

[38] Hosoki, A., et al.: Hetero-core structured fiber optic chemical sensor based on surface plasmon resonance using au/lipid films. Optics Communications 524, 128751 (2022).

[39] Zhang, J., Zeng, Z., Zhang, L., Lu, Q., Wang, K.: Application of mathematical morphological filtering to improve the resolution of chang’e-3 lunar penetrating radar data. Remote Sensing 11(5) (2019).

[40] Huang, W., Wang, R., Li, H., Chen, Y.: Unveiling the signals from extremely noisy microseismic data for high-resolution hydraulic fracturing monitoring. Sci. Rep. 7, 11996 (2017).

[41] Band, L.R., et al.: Root gravitropism is regulated by a transient lateral auxin gradient controlled by a tipping-point mechanism. Proceedings of the National Academy of Sciences 109(12), 4668–4673 (2012)

[42] Booth, A.D., et al.: Distributed acoustic sensing of seismic properties in a borehole drilled on a fast-flowing green-landic outlet glacier. Geophysical Research Letters 47(13), 2020–088148 (2020)

[43] Miyoshi, Y., et al.: Rice immediately adapts the dynamics of photosynthates translocation to roots in response to changes in soil water environment. Frontiers in Plant Science 13 (2023).

[44] Schindelin, J., et al.: Fiji: an open-source platform for biological-image analysis. Nat Methods 9(7), 676–82 (2012).

