## Supplementary Information for "Non-destructive real-time monitoring of underground root development with distributed fiber optic sensing"

Crop genetic engineering for better root systems can offer practical solutions for food security and carbon sequestration; however, soil layers prevent direct visualization. Here, we demonstrate an original device with a distributed fiber-optic sensor for fully automated, real-time monitoring of underground root development. We demonstrate that spatially encoding an optical fiber with a flexible and durable polymer film in a spiral pattern can significantly enhance sensor detection. After signal processing, the resulting device can detect the penetration of a submillimeter-diameter object in the soil, indicating more than a magnitude higher spatiotemporal resolution than previously reported with underground monitoring techniques. We also developed computational models to visualize the roots of root crops and monocotyledons, and then applied them to radish and rice to compare the results with those of X-ray computed tomography. The device's groundbreaking sensitivity and spatiotemporal resolution enable seamless and laborless phenotyping of root systems that are otherwise invisible underground.

### 1. SUPPLEMENTARY THEORY I

Here, we modeled the secondary root growth. Some dicotyledon crops, such as radishes and potatoes, store significant amounts of photosynthetic products in their roots during secondary growth. In agricultural practices, the size of each root determines the crop yield for these plants because a single root is directly consumed as a food source. For this purpose, we focused on the volume and modeled the lateral displacements of the soil as a single root increases its radius in a cylindrical system.

In this system, a dicotyledon was planted at the center of a cylinder filled with soil, where side wall consists of a PTFE film wrapped with the FOS, and the top and bottom bases were fixed. For simplicity, we assumed incompressible uniform soil, an intact PTFE film, and uniform radial growth by the root extending to the full depth of the cylinder.

Let the radii of the root be $r_0$ and the FOS be $r$. The original soil volume is $\pi r^2 h$ where $h$ is the depth of the cylinder. When the root volume $\pi r_0^2 h$ is introduced at the center, the film expands to $\sqrt{r^2 + r_0^2}$. Thus, the isodepth FOS deforms by $\frac{\Delta L}{L} = \frac{2\pi\sqrt{r^2+r_0^2} - 2\pi r}{2\pi r}$. Given that $\epsilon = \frac{\Delta L}{L}$ is known, the radius of the root can be determined as;

$$r_0 = r\sqrt{\epsilon^2 + 2\epsilon}. \tag{S1}$$

### 2. SUPPLEMENTARY THEORY II

Here, we modeled the primary root growth. For cereal plants such as rice and wheat, root system architecture is believed to determine crop yield. Thus, the extension of root growth is of interest. To determine the root coordinates in the soil, we focused on force propagation in the soil, in which the root tips act as force sources.

During primary root growth, the apical meristem divides and pushes the root cap downward to penetrate the soil. The force exerted by the root tips propagates through the soil and is sensed by the FOS. We assumed that the soil was homogeneous, isotropic, and linearly elastic in the full space . The point force solution for displacement $u$ in a full space is referred to as Kelvin's solution, and is given as

$$\mathbf{u} = \frac{1}{16\pi\mu(1-\nu)R}\left[(3-4\nu)\,\mathbf{P} + \frac{1}{R^2}\boldsymbol{\rho}(\boldsymbol{\rho}\cdot\mathbf{P})\right], \tag{S2}$$

where $\mu$ and $\nu$ are the shear modulus and Poisson ratio of soil, $R$ is the distance from the source to the sensor, $\mathbf{P}$ is the concentrated force at the point source, and $\boldsymbol{\rho}$ is the vector coordinates of the sensor. When a vertical stress $\mathbf{P} = P_z\,\mathbf{k}$ is applied at the origin, Eq. S2 in cylindrical coordinates are;

$$\begin{aligned}
u_r &= \frac{P_z}{16\pi\mu(1-\nu)R^3} rz, \\
u_\theta &= 0, \\
u_z &= \frac{P_z}{16\pi\mu(1-\nu)} \left[ \frac{(3-4\nu)}{R} + \frac{z^2}{R^3} \right],
\end{aligned} \tag{S3}$$

where $\boldsymbol{\rho} = \begin{bmatrix} r\cos\theta & r\sin\theta & z \end{bmatrix}^\mathsf{T}$ describes the cylindrical coordinate of the sensor and R = $\sqrt{r^2+z^2}$. While the strain tensor $\boldsymbol{\epsilon}$ is defined as

$$\boldsymbol{\epsilon} = 1/2(\nabla \mathbf{u} + \mathbf{u}\nabla), \tag{S4}$$

the base vectors in cylindrical coordinates are

$$\begin{aligned}
e_r &= \cos\theta i + \sin\theta j, \\
e_\theta &= -\sin\theta i + \cos\theta j, \\
e_z &= k,
\end{aligned}$$

and the partial derivatives in angular direction can be expressed as $\frac{\partial e_\theta}{\partial \theta} = -e_r$ and $\frac{\partial e_\theta}{\partial \theta} = -e_r$. Therefore, the strain components in Eq. S2 can be expressed as

$$\begin{aligned}
\epsilon_{rr} &= \frac{\partial u_r}{\partial r} \\
&= \frac{P_z}{16\pi\mu(1-\nu)R^3} z - 3\frac{r^2 z}{R^5} \\
\epsilon_{\theta\theta} &= \frac{u_r}{r} + \frac{\partial u_r}{\partial \theta} \\
&= \frac{P_z}{16\pi\mu(1-\nu)R^3} z \\
\epsilon_{zz} &= \frac{\partial u_z}{\partial z} \\
&= -\frac{P_z}{16\pi\mu(1-\nu)} \left[ (1-4\nu)\frac{z}{R^3} + 3\frac{z^3}{R^5} \right]
\end{aligned} \tag{S5}$$

The root coordinates were estimated using a hyperbolic line connecting the stem and root tip. Linearity and infinite space assumptions allow the superposition of point force solutions to represent multiple root tips.

### 3. SUPPLEMENTARY FIGURES

Fig. S1–S5 are supplementary figures.

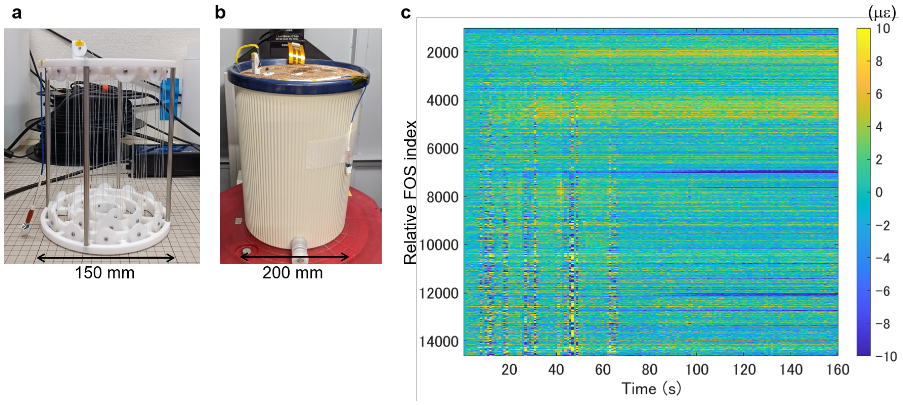


**Fig. S1.** Results of direct FOS installation in soil. (a) A photo of an early prototype composed of FOS directly supported by rigid structures. (b) The device installed in a large pot filled with Profile. (c) Distributed strain recording from the device when a manually directed metal wire with 1 mm-diameter penetrated into Profile at about 3 mm/s. Implementation of the direct FOS installation is described in Supplementary Methods.

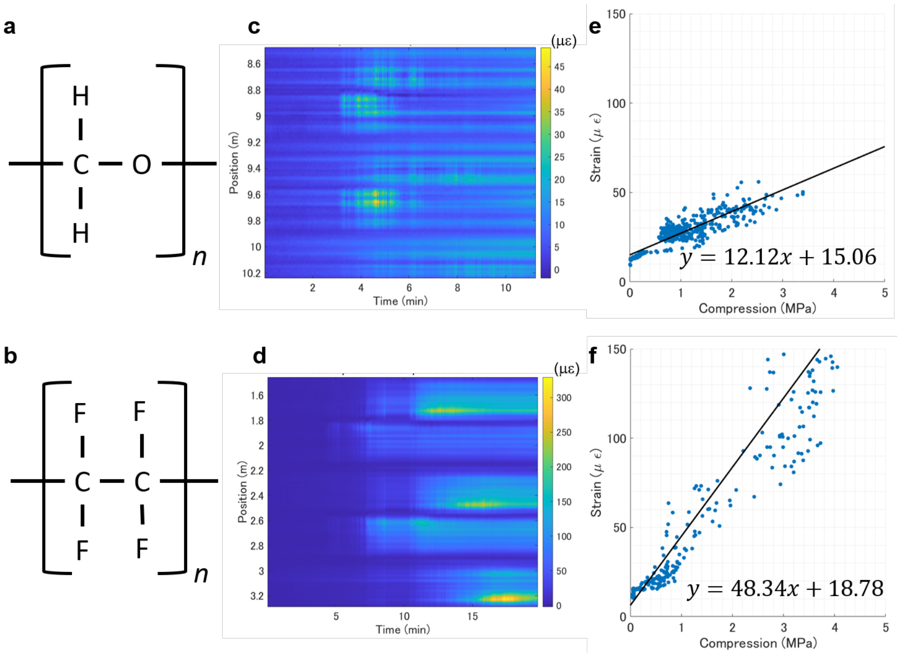


**Fig. S2.** Evaluation of FOS attached to polymer film. (a, b) Chemical formula for POM and PTFE. (c ,d) Distributed strain recording from the FOS attached to POM or PTFE film when 1 mm-diameter metal wire penetrated into Profile at 0.167 mm/s which filled the device and the surrounding. (e, f) Stress-strain curves of (c) and (d), respectively.

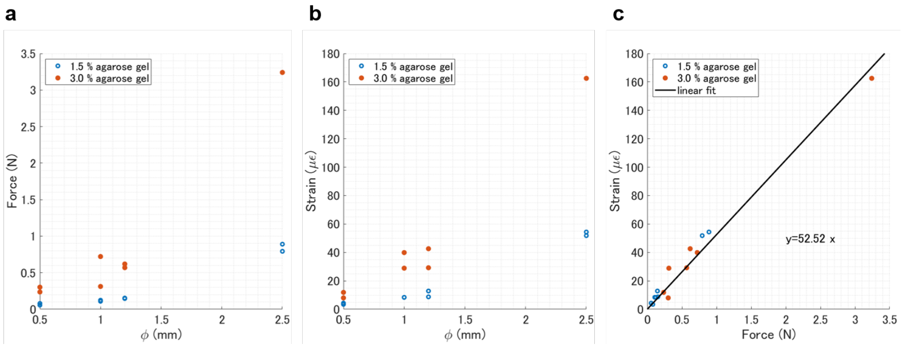


**Fig. S3.** Size, media stiffness, and force contributions to the strain response. (a) Scatter plot of reactive forces exerted by different agarose gel media upon penetrating cylindrical objects of various diameters. Maximal force is plotted for each penetration experiment. (b) Scatter plot of maximal strain detected by FOS in the experiments in a. (C) Scatter plot of the maximal strain versus maximal force in each experiment and the linear fit.

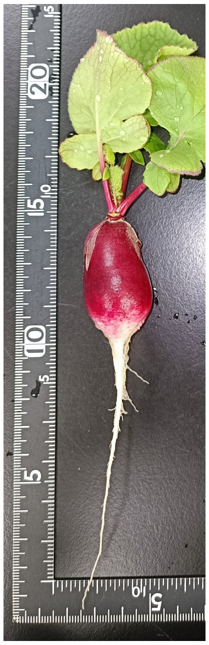

**Fig. S4.** A photo of the excavated radish.

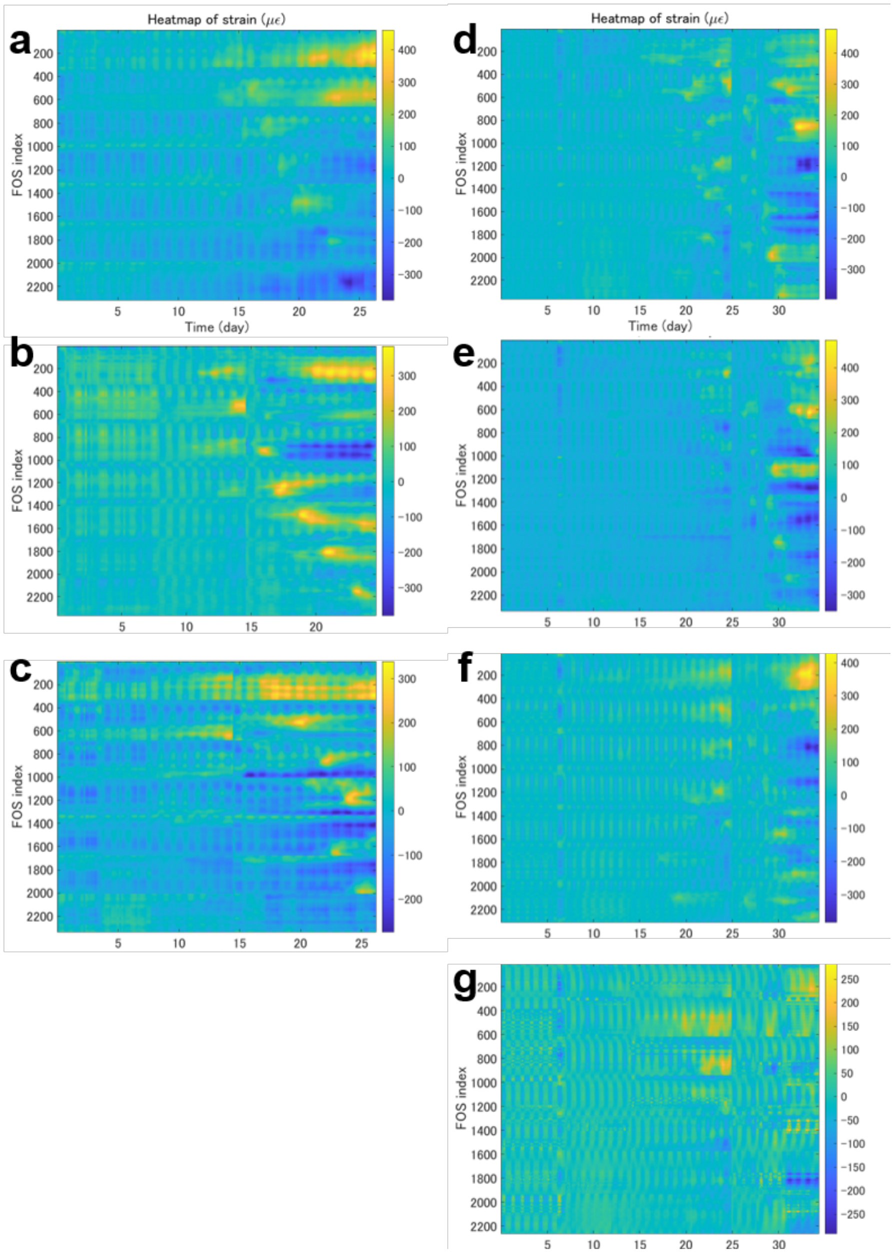


**Fig. S5.** Distributed strain recording from rice cultivation pots. Recordings were disrupted for the X-ray CT scans on day 15 and 27 for (a-c), and on day 14, 25, 28 for (d-g). (a) Data for Pot #1. (b) Data for Pot #2. (c) Data for Pot #3. (d) Data for Pot #4. (e) Data for Pot #5. (f) Data for Pot #6. (g) Data for Pot #7.

### SUPPLEMENTARY MOVIE

Supplementary movie of the radish root growth is provided externally.
